# Functional architecture of motion direction in the mouse superior colliculus

**DOI:** 10.1101/825711

**Authors:** Ya-tang Li, Zeynep Turan, Markus Meister

## Abstract

Motion vision is important in guiding animal behavior. Both the retina and the visual cortex process object motion in largely unbiased fashion: all directions are represented at all locations in the visual field. We investigate motion processing in the superior colliculus of the awake mouse, by optically recording neural responses across both hemispheres. Within the retinotopic map, one finds large regions of ∼500 μm size where neurons prefer the same direction of motion. This preference is maintained in depth to ∼350 μm. The scale of these patches, ∼30 deg of visual angle, is much coarser than the animal’s visual resolution. A global map of motion direction shows approximate symmetry between the left and right hemispheres and a net bias for upward-nasal motion in the upper visual field. Unlike other parts of the early visual system, the superior colliculus develops a locally biased representation of object motion.

## Introduction

The connection between the ecology of natural scenes and the neurobiology of vision has been a topic of enduring interest. To the first approximation, natural scenes are translation-invariant, meaning that any given visual feature or event is equally likely to happen anywhere in the image. This principle predicts that the neural representation of visual images should be approximately uniform everywhere in the visual field. Closer inspection suggests some departures from translation invariance. The upper visual field is often occupied by the sky, which differs in content from the lower visual field. Also animals direct their eyes to salient points in the scene, which selects for certain image features at the center of gaze.

The first station of the visual system seems to follow the principle of uniform coverage. In the mammalian retina, each of ∼30 distinct types of retinal ganglion cells (RGCs) conveys information about a specific visual feature^1, 2^. Each cell type tiles the entire retina in a “mosaic” arrangement, ensuring a representation of its special feature at every point in the visual space. The primary visual cortex continues this pattern. Individual neurons there are often tuned to one line orientation or motion direction, but all the possible orientations and directions are represented at every point, or at least within 1-4 degrees of visual angle^3–10^.

An evolutionarily older visual pathway leads from the retina to the superior colliculus^11^. In the superior colliculus, the principle of uniform coverage appears to be broken. Some of the earliest recordings in cat and rabbit already suggested a preference for visual motion in specific directions depending on location in the visual field^12, 13^. In mouse superior colliculus one finds large regions of the visual field, ∼30° across, where neurons prefer stimuli with a specific line orientation^14–17^. Whether the representation of visual motion is similarly biased in a coarse map remains debated^14, 18, 19^. Most of the studies to date were performed in anesthetized animals with an acute cortical lesion to uncover the underlying superior colliculus. Both of these perturbations are known to affect the direction-selectivity of neurons in the superior colliculus^17, 20, 21^.

The goal of the present work is to reveal how the direction of visual motion is represented in the superior colliculus of awake mice at a large-scale population level. To this end, we applied two complementary approaches. Two-photon calcium imaging was used to record neuronal responses at single-cell resolution in the posterior-medial SC of behaving mice, leaving the cortex intact. To reveal the functional organization on a larger scale over both hemispheres we applied a wide-field imaging method in a partial cortex mutant mouse.

## Results

### Single-cell imaging reveals a coarse map for direction of motion

To investigate the tuning of SC neurons to the direction of motion, we imaged neuronal calcium responses to dots and bars moving in different directions using two-photon microscopy in awake mice (Fig. 1a). To maintain the integrity of the overlying cortex, one is limited to the posterior-medial sector of the mouse SC that corresponds to the upper lateral visual field^15^ (Fig. 1b-c). Direction-selective (DS) SC neurons in this region responded robustly to both moving dots and moving bars. Many neurons had a strong preference for one direction (Fig. 1d). We defined an index of direction selectivity (DSI, see Methods) and focused further analysis on neurons with DSI larger than 0.1 (Fig. 1e-f). Among these cells, the response to preferred direction motion was ∼3 times larger on average than that to the null direction.

**Figure 1.**
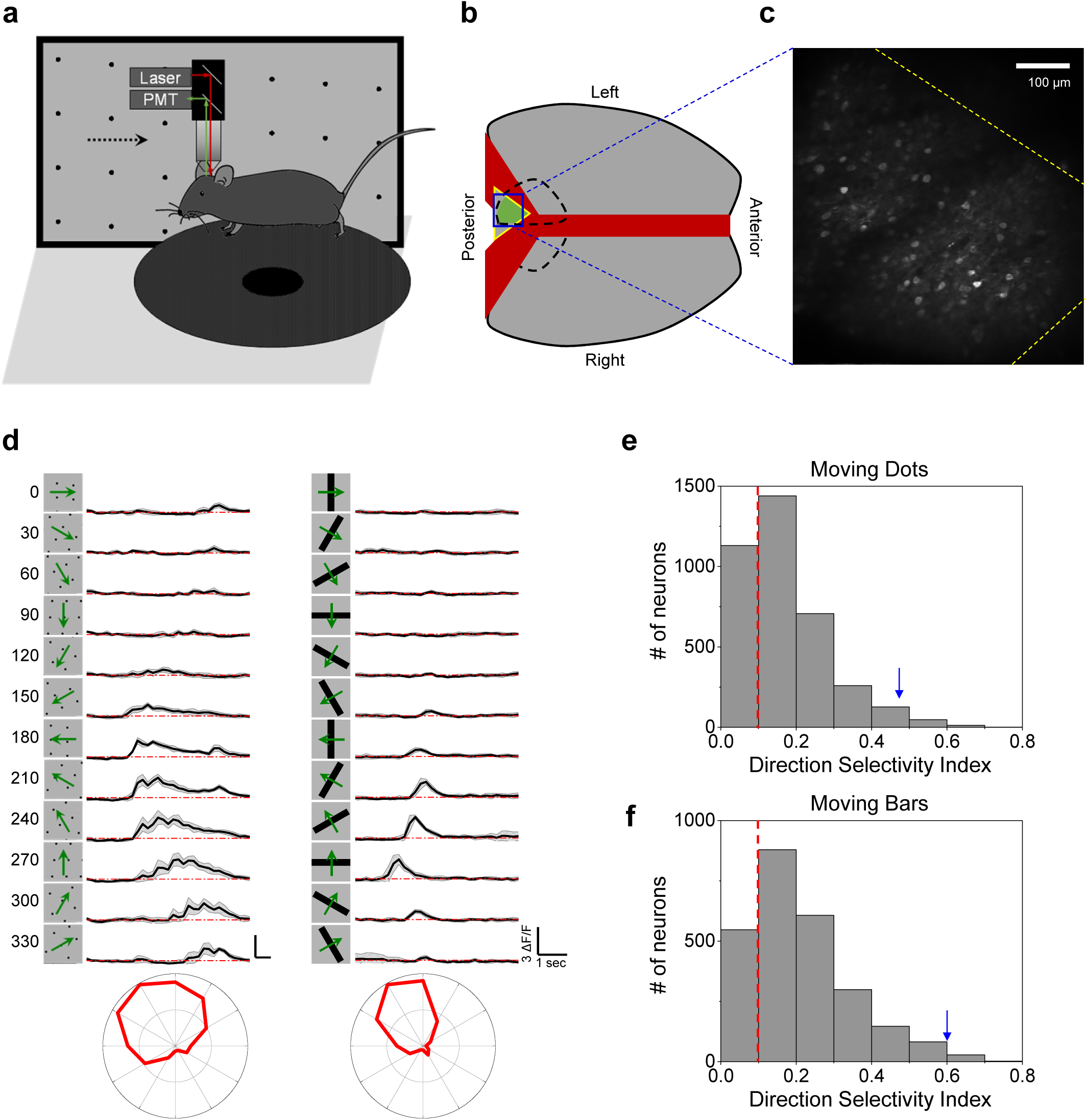
Two-photon calcium imaging reveals direction tuning in awake mouse superior colliculus. **a**, Schematic of the experimental setup. Mice were head-fixed and free to run on a circular treadmill. Visual stimuli were presented on a screen. Neuronal calcium activity was imaged using two-photon microscopy. PMT, photomultiplier tube**. b**, Schematic of mouse brain anatomy after insertion of a triangular transparent plug to reveal the posterior-middle part of the SC. **c**, A standard deviation projection of calcium responses to moving dots. **d**, Calcium responses of a sample neuron to moving dots and moving bars at twelve directions. Gray shade indicates the standard deviation across identical trials. Small wiggles in the moving dot response reflect individual dots crossing the receptive field. Bottom, polar graphs of normalized peak responses. **e**, Histogram of direction selectivity index of neurons responding to moving dots. Dashed line indicates the threshold applied for direction selectivity analysis. Arrowhead marks the cell in (d). **f**, Histogram of direction selectivity index of neurons responding to moving bars, displayed as in (e).

The moving-bar stimulus will drive feature detectors that are tuned to either motion direction or line orientation, leading to possible confounds, whereas a field of moving dots has no orientation signal. Therefore, we focused on results from moving dot stimuli. Figure 2 illustrates the direction tuning of neurons in the left posterior-medial SC of three different animals. In each case, neurons located near each other tended to have the same preferred direction of motion. That preferred direction varied only gradually across the surface of the SC (Fig. 2a-d), or remained constant over distances of several 100 µm (Figs 2e-h, 2i-l). At times, neurons that prefer two opposite directions were intermingled (Fig. 2i-l). None of these arrangements of preferred direction resembles the salt-and-pepper organization in rodent primary visual cortex (V1)^10^ (see more examples in Supplementary Fig. 1). Instead, the preferred direction changes only slowly with the location on the SC, if at all. The regions of constant direction bias range up to ∼500 μm in size, which corresponds to ∼40 degrees of visual angle or 4 receptive-field diameters^17^. Furthermore, in all 3 animals, the preferred direction is the same, suggesting that the posterior-medial SC has a systematic bias for upward-nasal image motion.

**Figure 2.**
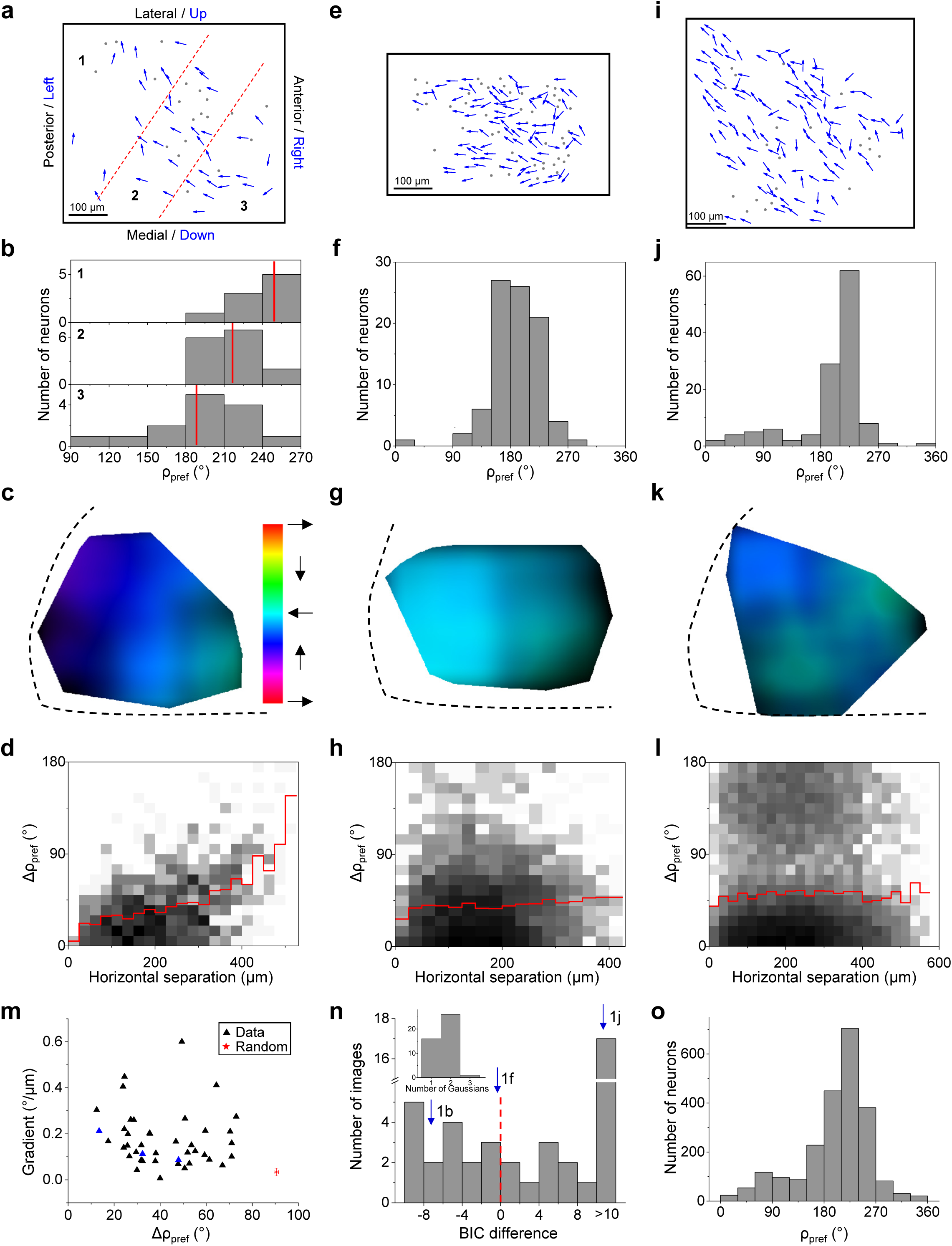
Locally uniform direction tuning. **a**, All direction-selective neurons in a sample field of view from left SC. Arrows are plotted at the anatomical location of the neurons and indicate the preferred direction in monitor coordinates (blue axis legend). Dots are neurons with low direction-selectivity. Dashed lines divide the image into three regions. **b**, Histograms of preferred direction for the three regions. Red lines mark the mean. **c**, A smoothed version of the direction map in (a) where the hue encodes the preferred direction and the intensity varies inversely with the local variance of preferred directions. Note the systematic color gradient across the image. Spatial scale as in (a). **d**, A 2-D histogram of the absolute difference in preferred directions of two neurons versus their horizontal distance (41 cells, 820 pairs). The red line indicates the mean. (**e**, **f**, **g**, **h**) and (**i**, **j**, **k**, **l**) are single fields of view from two different mice represented in the same way as (**a**, **b**, **c**, **d**) with (88 cells, 382 pairs) and (124 cells, 7626 pairs) respectively. The dark regions in **k** indicate large local variance. **m**, The gradient of the map plotted against the absolute difference in preferred directions for pairs within 50 µm for all fields of view (42 images from 14 animals). Blue triangles are from the three sample images in (c), (g), (k). Red star indicates the mean of 1000 samples from a simulated random arrangement of preferred directions; error bar indicates SD. **n**, Histogram of the difference of Bayesian information criterion (BIC) between one-Gaussian and two-Gaussian fitting for all fields of view (42 images from 14 animals). Dashed line indicates the boundary between unimodal and bimodal. Arrows mark the three sample images. Inset, the number of Gaussians that gives the minimal BIC. **o**, Histograms of preferred direction for all direction-selective neurons (n = 2279 cells from 14 animals).

To summarize the results across many recordings, we calculated for each field of neurons both a measure of the local scatter of preferred directions as well as the systematic gradient in the spatial map of preferred direction (Fig. 2m). The resulting distribution diverged dramatically from the null model in which each cell has an independent preferred direction (Fig. 2m). In more than half of the cases, the preferred direction showed a bimodal distribution (Fig. 2n). Across all samples, the preferred direction was strongly biased towards ∼220° (upward/nasal) (Fig. 2o). This biased representation of motion direction was also observed in the responses to moving bars (Supplementary Fig. 2).

To explore the organization of preferred direction in the vertical dimension, we imaged neuronal responses up to ∼350 µm deep in steps of 20-30 µm. We projected the direction maps from different depths onto the same plane (Fig. 3a). This revealed that neurons with short horizontal separation preferred a similar direction independent of depth (Fig. 3a-b). Further analysis over 11 animals confirmed that preferred direction has a columnar organization, in that it varies little in the vertical dimension (Fig. 3c).

**Figure 3.**
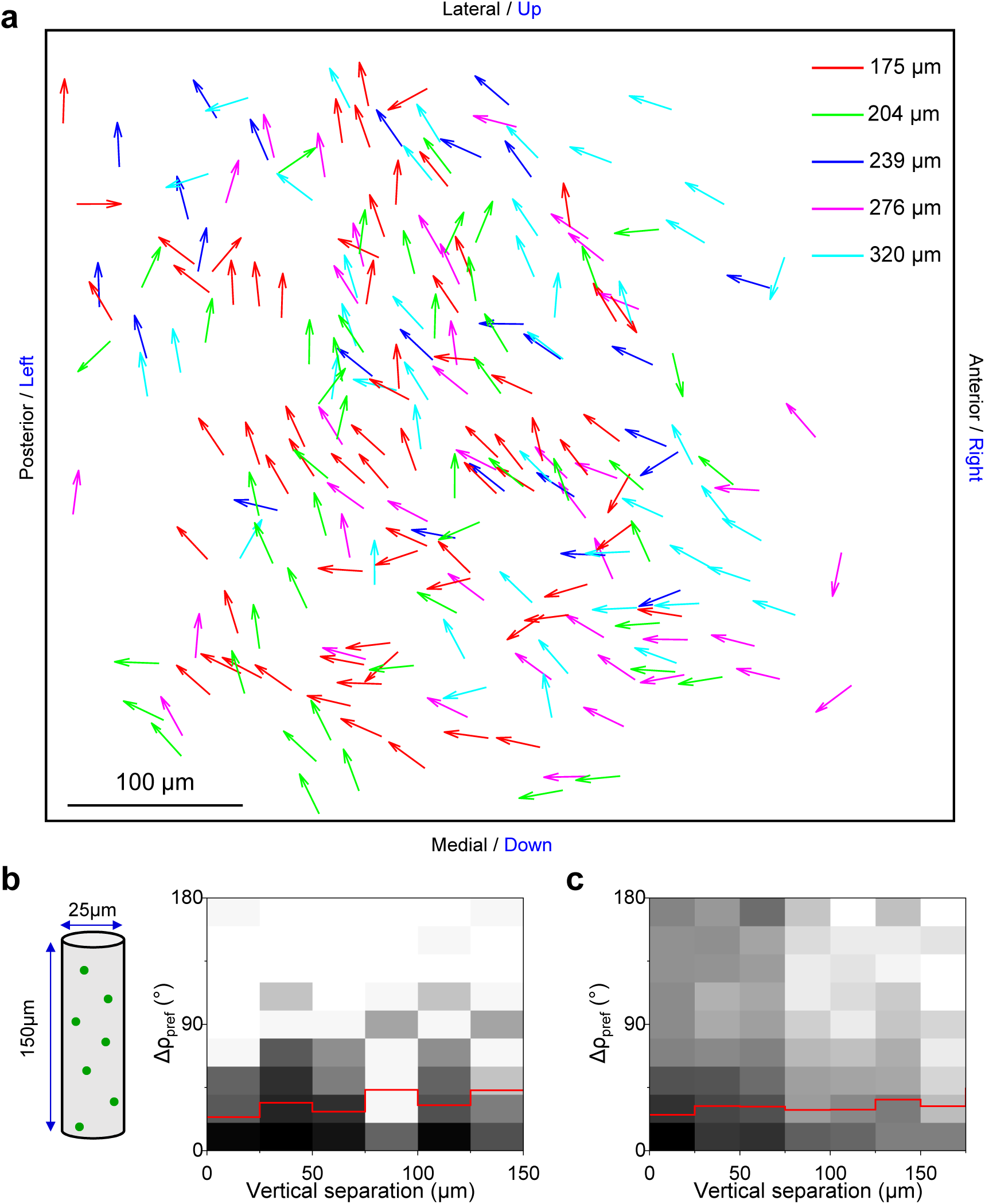
Direction columns extend in depth. **a**, Horizontal plane projection of arrow plots (as in Fig. 2a) at five different depths in the same field of view (229 cells). **b**, 2-D histogram of the absolute difference in the preferred direction versus vertical separation for this volume (236 pairs). Red line indicates mean. **c**, 2-D histogram of the absolute difference in the preferred direction versus vertical separation for all recorded neurons (2323 pairs from 11 animals).

### The preferred direction is orthogonal to the preferred orientation

Early work on monkey and cat visual cortex discovered an orthogonal relationship between preferred orientation and preferred direction^6, 22^. We examine whether this claim can be generalized to the mouse SC. To probe these two maps independently, we measured the direction preference with moving dots that carry no orientation signal and the orientation preference with flashed gratings that carry no motion signal (Fig. 4a). Analysis of direction- and orientation-selective neurons in an entire window of cells spanning 580 μm revealed that the angle between their preferred direction and preferred orientation is about 90° (Fig. 4b-c, Supplementary Fig. 3a-b). The orthogonal relationship was confirmed by histogramming the difference between preferred direction and orientation for neurons across many animals (Fig. 4d).

**Figure 4.**
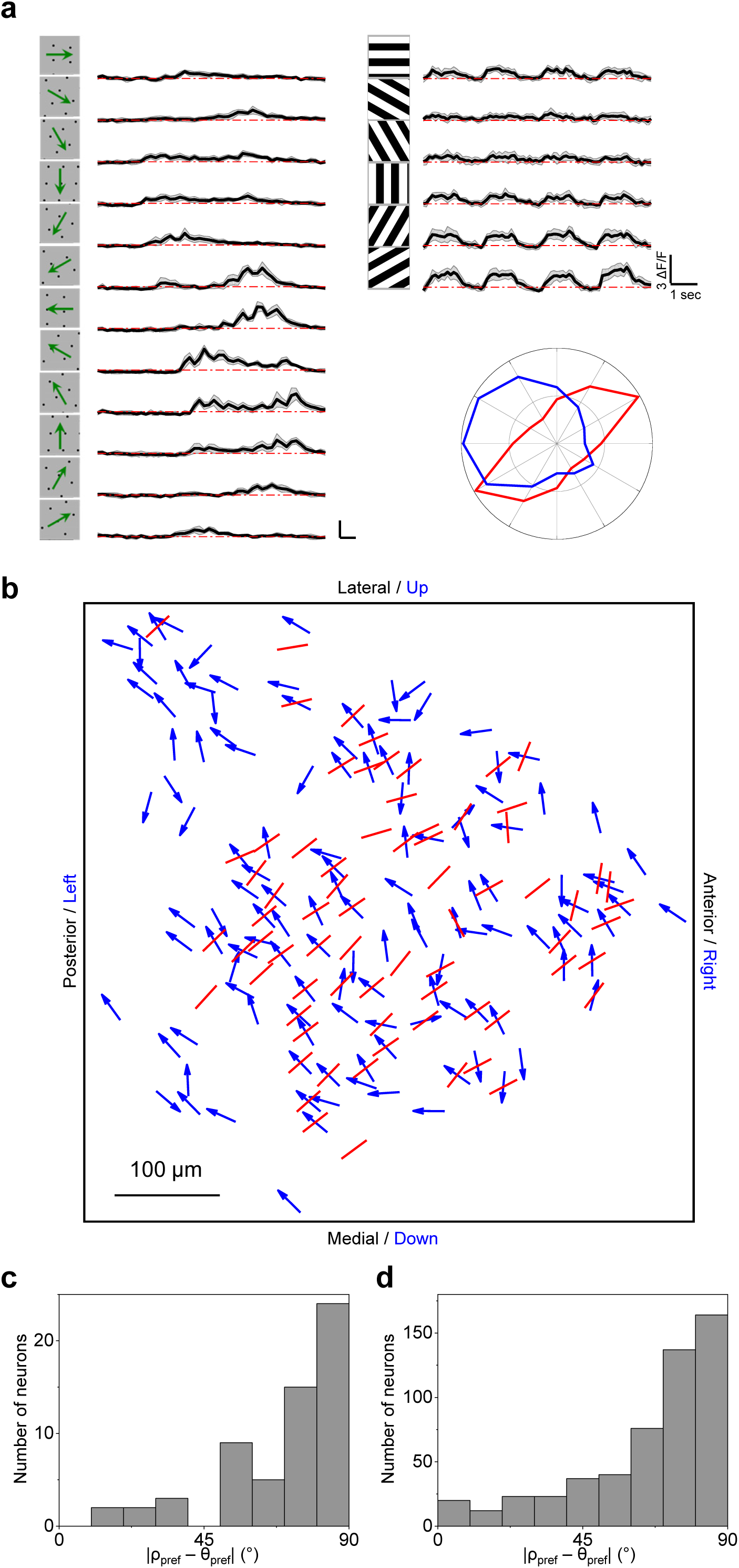
Direction maps are orthogonal to orientation maps. **a**, Calcium responses to moving dots and flashed gratings for a sample neuron exhibiting both direction- and orientation-selectivity. Polar plot shows peak response to dots (blue) and gratings (red). Here the preferred direction and orientation differ by 86°. **b**, Overlay of maps for preferred direction (arrows) and orientation (lines) in a sample field of view, displayed as in Fig. 2a. **c**, Histogram of the absolute difference between preferred direction and preferred orientation for the sample in (b) (60 cells). **d**, Same histogram for all direction- and orientation-selective neurons (532 cells from 14 animals).

### Wide-field imaging reveals the direction maps of two entire hemispheres

The bias in the preferred direction towards upward motion among neurons in the posterior-medial SC (see Fig. 2o) motivated us to explore the functional organization of motion direction on a larger scale. To this end, we recorded from the entire SC using a wide-field imaging technique that sacrifices single-cell resolution but promised a global view of the gradual variations in direction tuning. We first validated the approach by imaging responses in the same brain using both two-photon and wide-field microscopy. Figure 5a-b illustrates the direction map in the posterior-medial sector of the SC obtained by both techniques, with very similar results. For broader coverage, we imaged the SC of mutant mice whose posterior cortex fails to develop^23^, leaving almost the entire SC directly accessible for optical recording (Supplementary Fig. 4). By repeatedly moving the narrow window of the 2-photon microscope, we accumulated single-cell recordings of the preferred motion direction for ∼600 neurons over much of the left SC (Fig. 5c-d). A subsequent repeat of the same stimuli using the wide-field microscope yielded essentially the same map of preferred direction (Fig. 5e). Furthermore, the direction map in the posterior-medial segment is the same in partial cortex mutant and normal animals (compare to Fig. 5a-b). These results show that wide-field imaging is a valid approach to reveal the functional organization of direction preference at a large scale.

**Figure 5.**
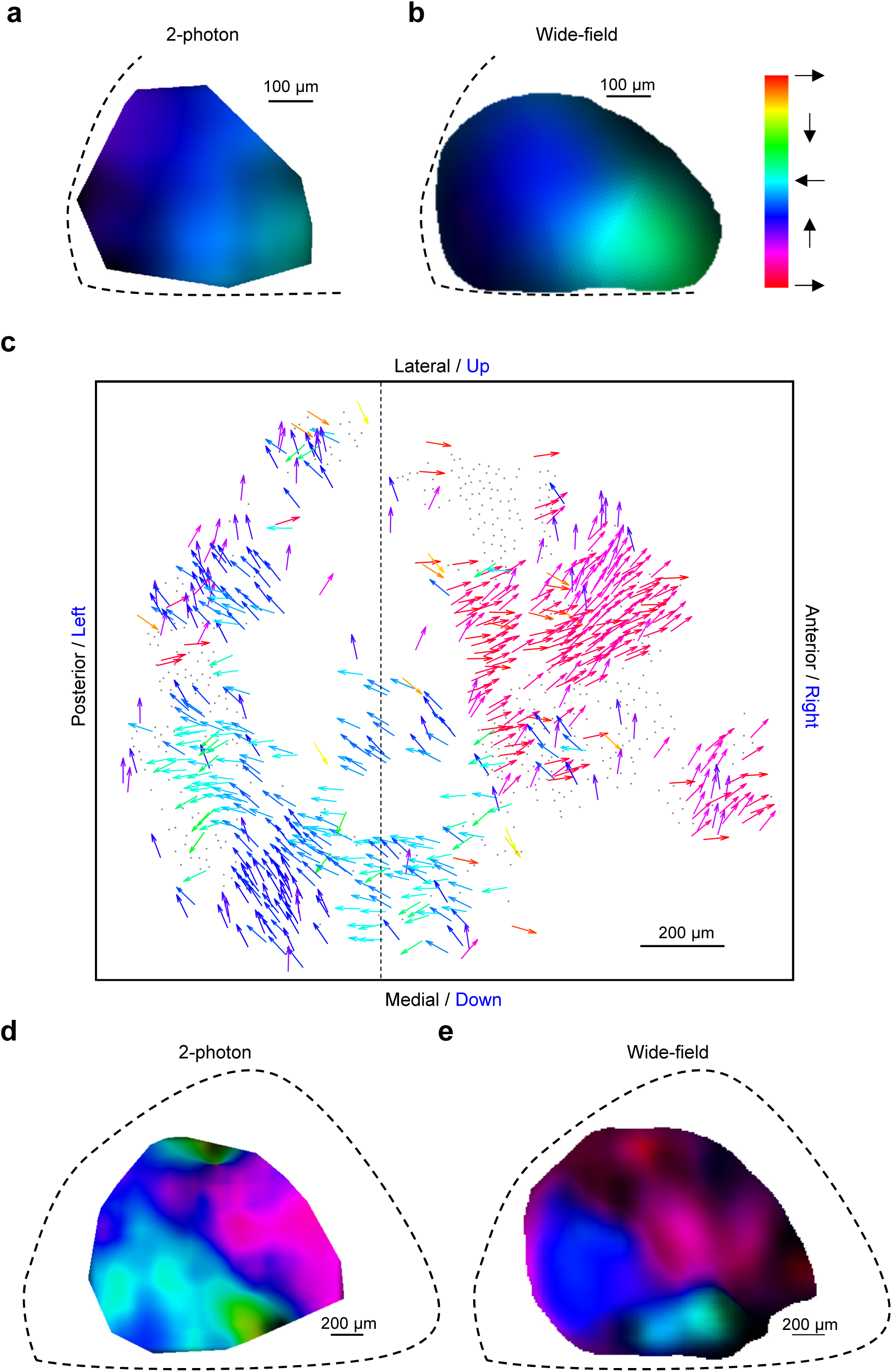
Direction maps from wide-field imaging and two-photon imaging agree. **a and b**, Direction maps generated from two-photon data and wide-field imaging data in the posterior-medial left superior colliculus of the same mouse. Hue represents the preferred direction (a) and value codes the local variance (a) or the DSI (b). **c**, All direction-selective neurons in the SC of a mutant mouse with undeveloped cortex presented in the anatomical space with the preferred direction in monitor coordinates as in Fig. 1a. Dashed line separates images that were acquired with the monitor at two different azimuth coordinates: 95° for the left and 45° for the right. **d**, A smooth direction map from interpolating the data in (c). **e**, Direction map from wide-field imaging data from the same animal as in (d).

The patchy anatomy of preferred direction on the SC suggests that large parts of the visual field experience a bias for certain motion directions. To investigate this functional map, one needs to represent each neuron at the location of its receptive field, along with its direction preference in visual field coordinates. The need to represent motion directions throughout the entire visual field brings up an interesting challenge for graphic display. The conventional spherical coordinate system for visual space has singularities at the “north” and “south” poles (elevation = +90° and −90°) where the direction of motion cannot be expressed with respect to the coordinates (at the North Pole all directions point south). But the north pole of the spherical coordinate system is well within the visual field of the mouse^24^. To eliminate these pathologies, we chose to represent the sphere with a “unipolar” coordinate system that has only one singularity. That special point can be placed dead behind the animal, outside of the visual field (Fig. 6a, Supplementary Fig. 5, see **Methods**). In the following, we will use these unipolar coordinates for large-scale maps of the mouse visual field.

**Figure 6.**
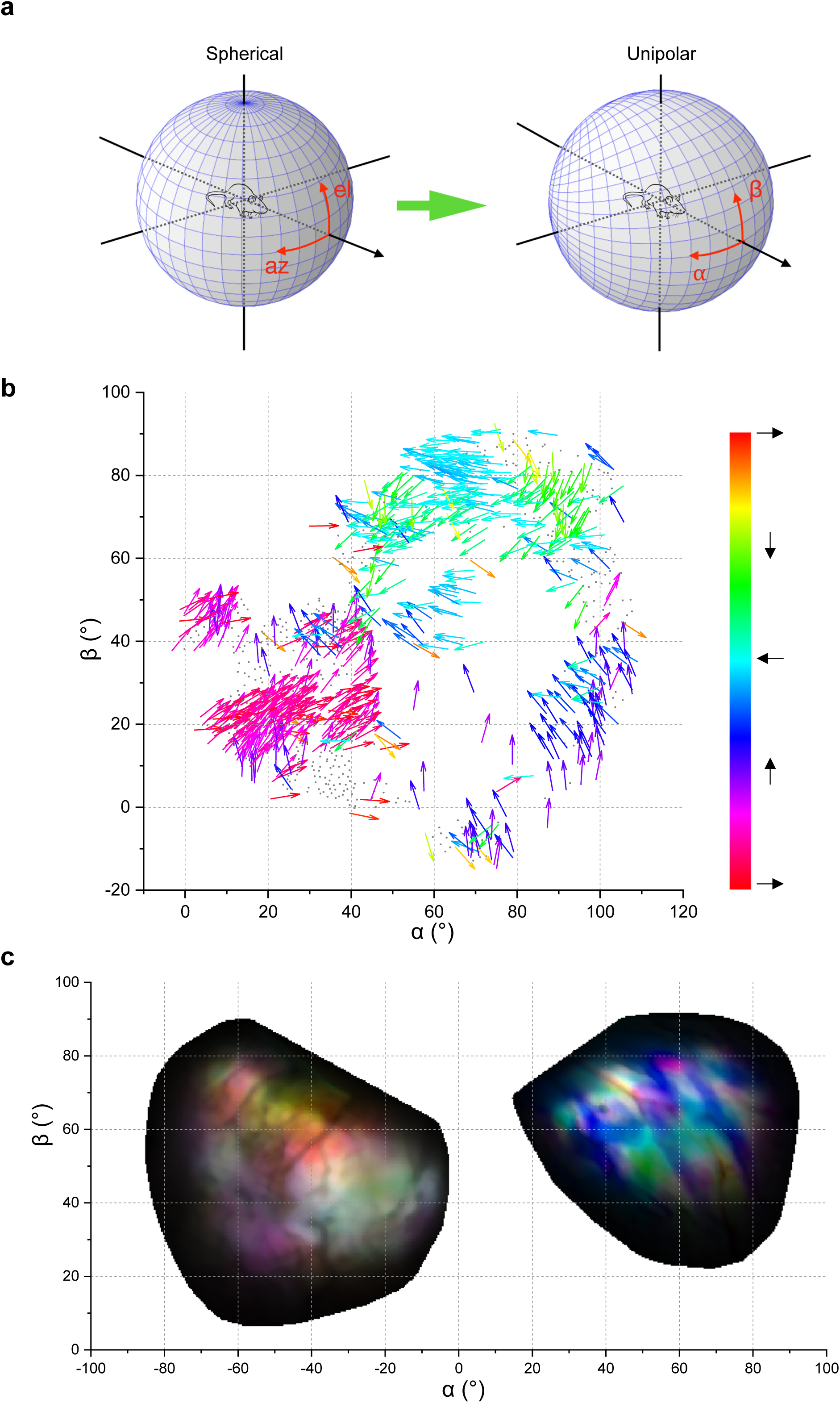
Large-scale direction maps in visual space. **a**, Schematic illustrating the correspondence of spherical coordinates and unipolar coordinates. **b**, All direction-selective neurons in the SC of a partial cortex mouse presented in the visual space using unipolar coordinates. Same recording as Fig. 5c, but the arrows are plotted at receptive field locations. **c**, Direction map obtained from both left and right SC of a different mouse, plotted in visual space with unipolar coordinates. Hue codes the preferred direction; value codes the amplitude of responses; saturation codes the DSI.

For the sample recording of Fig. 5c we mapped the receptive fields and preferred directions of all neurons into visual space (Fig. 6b). As predicted, the preferred direction varies with location in the visual field. Patches of approximately constant bias, ranging from 10° to 40° in size, are separated by regions with sharp changes. The gradient of preferred direction varies across the visual space. Wide-field imaging performed across both sides of the brain confirmed the patchy organization and the net bias for upward motion. Furthermore, neurons representing the left and right visual fields appear to prefer mirror-symmetric directions (Fig. 6c, Supplementary Fig. 6).

To identify the systematic aspects of direction-sensitivity, we analyzed these global maps across many animals. The simplest overarching feature is the overall bias for certain directions of motion, slightly nasal of upward, which is mirror-symmetric across the two sides of the visual field (Fig. 7a). To quantify the size of the patches of constant direction, we calculated the autocorrelation function for each map, which yielded a full width at half maximum of 30° (Fig. 7b, Supplementary Fig. 7a). As might be expected this average patch size is the same in the left and right half of the SC (Fig. 7c). The direction patterns in left and right visual fields do not align perfectly: A cross-correlation between the two sides has a somewhat wider peak of ∼40° (Fig. 7c). This concordance is similar to that observed across different animals (Fig. 7c, Supplementary Fig. 7b-c). Figure 2 illustrates the variation across animals at single-cell resolution.

**Figure 7.**
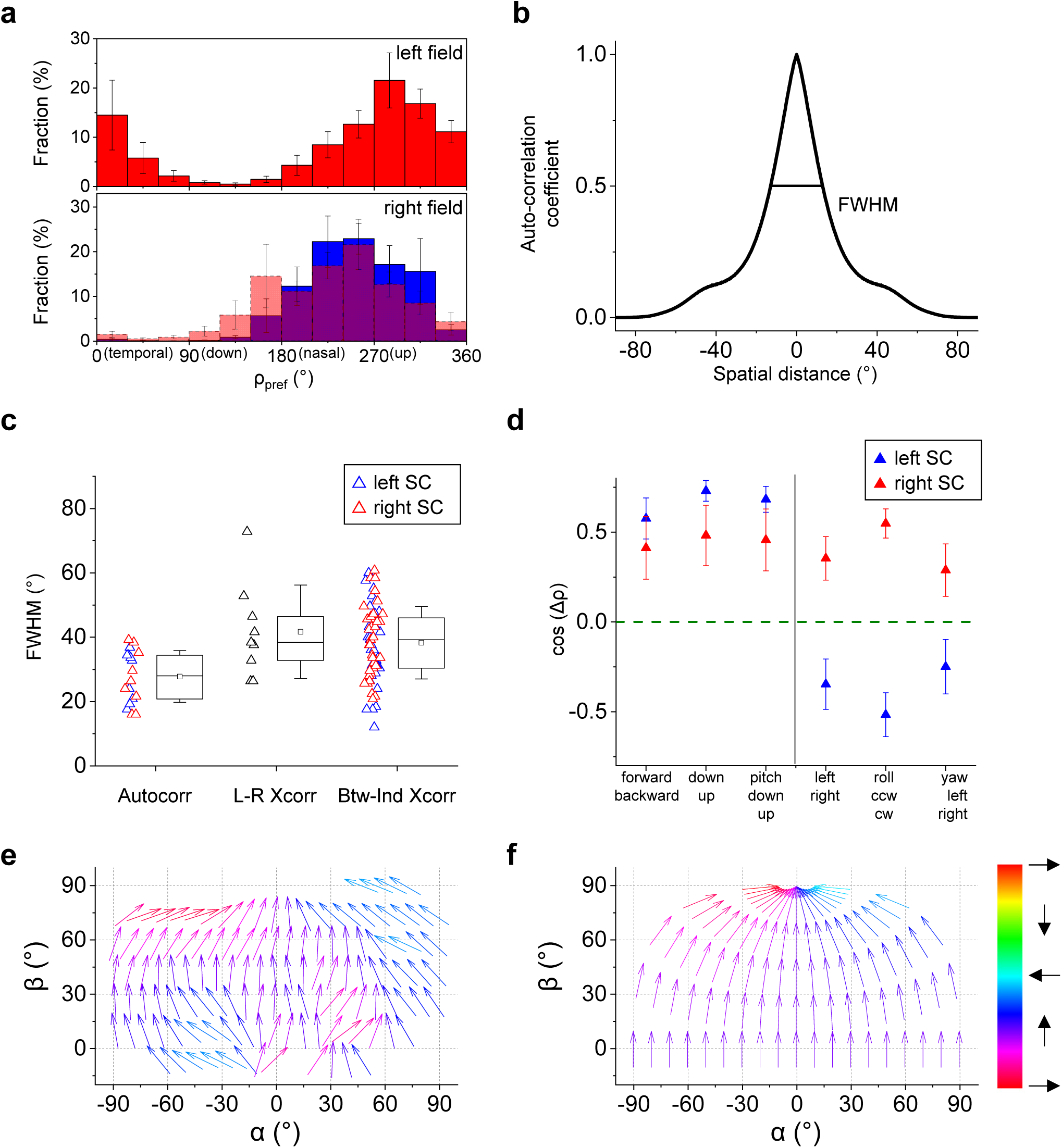
Direction bias in the direction map. **a**, Distribution of preferred directions encountered in both visual hemifields (9 animals). Error bars represent one standard error across animals. Bottom: The histogram for the left hemifield was flipped about the vertical and overlaid on the histogram for the right hemifield. **b**, The radial profile of autocorrelation for a sample direction map. Full-width at half maximum (FWHM) is indicated. **c**, Box plots of the FWHM for the autocorrelation, the cross-correlation between left and right SCs, and the cross-correlation between individuals for the same side of the SC (9 animals). The box plots 25^th^ to 75^th^ quartiles, the band inside the box is the median, and the square inside the box is the mean. The ends of the whisker indicate one standard deviation above and below the mean. **d**, Projections of preferred-direction maps from experimental data onto motion maps representing different optic flows (9 animals). Error bars represent one standard error. Wilcoxon signed-rank test or one-sample *t*-test were applied to test the significance of the difference from 0; the p-values are 1.5e-3, 4.3e-6, 3.1e-5, 0.043, 3.9e-3, and 0.14 for the left SC; 0.098, 0.039, 0.027, 0.019, 1.5e-4, and 0.08 for the right SC. Mann-Whitney U-test or two-sample *t*-test were applied to test the significance of the difference between left and right SCs, and the p-values are 1.8e-3, 2.2e-6, and 0.022 for the right half figure. **e**, Arrow plots of the preferred direction in the SC, averaged over 9 animals, and plotted across the visual field in unipolar coordinates. **f**, The motion vectors in the mouse’s visual field produced by an aerial predator approaching at constant height from various directions.

One possible interpretation of the global inhomogeneity is that it relates somehow to the optic flow pattern created by self-motion of the animal. To evaluate this notion, we compared the experimental preferred-direction maps to optic flow maps predicted from the six basic modes of self-motion (Supplementary Fig. 8). By projecting the optic flow map onto the direction map one gains an estimate of how strongly the stimulus activates the superior colliculus. For those self-motions without a lateral component, the two sides of the SC are tuned in the same way. In particular, they are more strongly activated by forward translation, down translation, and downward pitch (Fig. 7d). For those motions with a lateral component, the two sides are tuned in the opposite way. In particular, the left SC prefers translation to the right, clockwise roll, and yaw to the right; *vice versa* for the right SC. Most of these biases are highly significant and consistent across animals.

Alternatively one can relate the motion bias of the superior colliculus to movements in the visual scene outside of the animal. In the upper visual field, there is an intriguing match between the direction tuning of the SC and the motion vectors produced by an aerial predator approaching at a constant height from the horizon: Such a bird’s trajectory describes an upward path in the visual field that eventually bends in a nasal direction towards the north pole (Fig. 7e,f).

## Discussion

This study extends our understanding of the functional architecture of visual features in the brain. We showed that the mouse superior colliculus represents visual motion with a bias that depends on location in the visual field. Nearby neurons prefer a similar direction of motion even over distances of ∼500 μm on the collicular surface, with an average patch size of ∼30 degrees of visual angle (Fig. 2). The direction bias extends vertically over several 100 µm to form columns of preferred direction (Fig. 3). For neurons that are selective both for direction and orientation, the preferred direction is orthogonal to the preferred orientation (Fig. 4). Large-scale wide-field imaging revealed a global map of preferred direction with coarse symmetry between two sides of the superior colliculus and a strong bias for upward-nasal motion in the upper visual field (Figs. 5–7). Here, we briefly discuss these conclusions and their implications.

### Relation to prior work

Some of the features reported here have been described before: For example, de Malmazet et al^18^ report that nearby neurons in certain regions of the retinotopic map tend to prefer the same direction of motion, and that this local preference also extends in depth. There has been controversy on those points^17, 19^, but our results reliably confirm the observation of a local direction bias. Building on this prior work, we here present a global analysis of direction coding in the SC. We extend the measurements to a larger visual field, especially in the superior elevations ignored in previous studies. We also record from both sides of the brain, covering almost the entire surface of the SC accessible from above. We established that the global map consists of multiple iso-direction patches and measured their size. Furthermore, we assessed the symmetry of these patches across the two sides of the SC and evaluated how the global direction map interacts with the global optic flow fields that result from locomotion during natural behavior.

Our results differ from prior claims on the topic of orientation tuning and how it relates to direction tuning. The present observation that preferred orientation systematically lies perpendicular to the preferred direction is consistent with the relationship reported in neurons of the visual cortex^5, 6, 22^. This contrasts with a prior report^18^ which argued that direction maps and orientation maps in mouse superior colliculus are independent. That conclusion may have been affected by the choice of stimuli, which precluded an independent measurement of orientation and direction tuning in the same neuron (Supplementary Fig. 9). In the present study, tuning to line orientation vs object motion was assessed by different stimulus probes.

### Functional anatomy in different species

The functional organization of visual processing has been reported in the brains of many species. The visual cortex of cat^10^, monkey^6^, and ferret^5^ shows a functional map, where a neuron’s response features – such as preferred orientation and direction – depend systematically on its location in the brain. In the rodent cortex there is less organization, and neurons preferring different orientations are intermingled in a “salt-and-pepper” pattern. Despite this difference, the common point is that neurons of all preferred orientations (or directions) are represented within a visual region smaller than the receptive field of a single neuron. In the primary visual cortex (V1) of the monkey, the orientation columns measure about 0.1° on average, only one-tenth of the receptive field size^25^. In V1 of the tree shrew, all orientations are represented within a 4° visual angle, again smaller than the average receptive field size^9^. By contrast, in the mouse superior colliculus, the patches with uniform preferred orientation span ∼30°, which greatly exceeds the RF size of ∼10°^17^, or the visual acuity of 2°^15^. The results reported here show that the representation of object motion in the superior colliculus is similarly dominated by such local biases. An intriguing new feature is that the iso-direction patches contain a minority of neurons tuned to the diametrically opposite direction (Fig. 2i,j,k,l). It will be important to understand the relationship of these two neural populations within the local circuitry.

### How does the patchy direction map in the SC arise?

Several explanations can be contemplated: (1) The local direction bias may exist already in the neural projections from the retina. (2) It may result from selective connectivity between the retinal inputs and the target neurons in the SC. (3) Or it may be constructed by postsynaptic circuitry within the SC. It has been shown that much of the direction selectivity in the mouse SC arises already in the excitatory synaptic inputs from retinal ganglion cells^26^, which argues for a diminished role of collicular interneurons in the phenomenon. The mouse retina contains 8 types of direction-selective ganglion cells (DSGCs). Their global distribution across the retina has been analyzed, and there is no indication of local inhomogeneities that could explain the patchy organization in the direction map of the SC^27–29^. Furthermore, the projections of DSGCs to the superficial SC appear to form a layer of uniform density, with no indication that one subtype dominates over the others in any local region^30, 31^. This prior work suggests that each retinotopic location on the SC receives an equally balanced input of direction-selective signals from the retina, quite unlike what occurs in the optic tectum of the zebrafish^32^. If so, then the local bias observed in iso-direction patches must result from selective connectivity between SC neurons and specific subtypes of DSGCs, and that connectivity bias must somehow be coordinated among nearby neurons. Conceivably the local circuitry of interneurons contributes to this lateral coordination.

### Behavioral relevance

Is there an intelligible purpose to the direction architecture in the mouse superior colliculus? On a coarse scale, cells in the upper visual field systematically prefer motion in the upward and nasal direction. Because activity in the upper visual field is generally associated with defensive reactions^33, 34^, one may speculate that the direction tuning is adapted to those functions. Indeed, we found that the global pattern of preferred direction across the visual field aligns with the motion vectors that would be produced by an aerial predator approaching from the horizon.

In addition, the global bias in the upper visual field also predicts a selectivity for certain optic flows produced by self-motion. For example, the superior colliculus as a whole should respond more strongly to forward motion of the animal than to backward motion, whereas horizontal turns left or right should produce excess activity in the right and left colliculus respectively. Such strong global signals may play a role in rapid feedback during the control of locomotion.

The other characteristic scale of the direction architecture is the size of smaller patches that deviate from the average tuning, measuring ∼30° of visual angle. The meaning of these specializations is harder to discern. The effects are substantial: For a direction-selective neuron in such a patch, the median ratio between preferred and null direction response is about 2.2. Thus, differently tuned patches may differ by a factor of 4 in their relative response to two moving objects. This should produce noticeable regional gradients in the animal’s perceptual sensitivity to motion, a prediction to be tested in future experiments.

## Methods

### Visual field coordinate systems

During experiments with visual stimulation, the animal was head-fixed with the lambda-bregma plane horizontal in the laboratory frame. To report locations in the visual field, we use either spherical coordinates (azimuth,elevation) = (*φ,θ*) or unipolar coordinates (*α,β*). The lambda-bregma plane is at *θ* = 0° and *β* = 0°.

### Unipolar Coordinates

We introduce this coordinate system to avoid the singularities occurring at the poles of spherical coordinates. Suppose the observer sits at the origin of a Cartesian system facing in the *z*-direction (Fig. 6a). The horizontal plane is (*x*, *z*) and the *y*-axis points up. A unit sphere is centered on the origin. For a given point 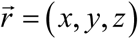 on the sphere, spherical coordinates (azimuth,elevation) = (*φ,θ*) are defined as

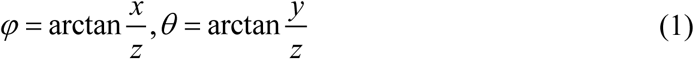

and unipolar coordinates (*α,β*) as

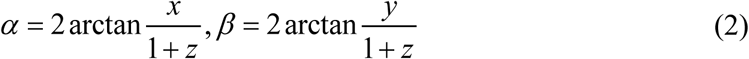

Note the only singular point of the unipolar system is directly behind the observer at 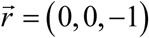. The transformation from spherical coordinates to unipolar coordinates is:

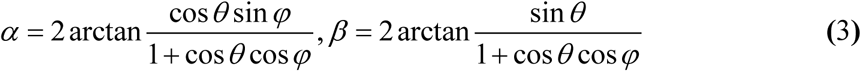

The direction of motion on the sphere in unipolar coordinates is defined as

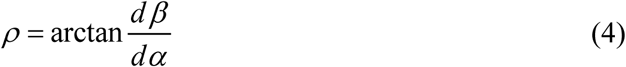

where *dα* and *d β* are increments of motion along *α* and *β* respectively.

### Animal subjects

Laboratory mice of both sexes were used at age 2-4 months. The strains were C57BL/6J (wild-type) and Emx1-Cre:Pals1^flox/wt^ (partial cortex^23^). All animal procedures were performed according to relevant guidelines and approved by the Caltech IACUC.

### Visual stimulation

For 2-photon imaging experiments, an LED screen was placed 15 cm away from the mouse’s right eye. The center of the monitor was at 95° azimuth and 25° elevation in the laboratory frame, and the monitor covered a visual field of 106° x 79°. The monitor was strobed for 12 µs at the end of each laser scan line to minimize interference of the stimulus with fluorescence detection. For measuring neuronal tuning to orientation and motion direction we applied three types of stimuli. (1) A 2-d array of moving dots with a diameter of 5° spaced at 54°. The array moved in 12 possible directions. This stimulus contains no orientation signal. (2) Square gratings with a spatial frequency of 0.08 cycles per degree were flashed at 6 orientations and 4 phases. This stimulus contains no motion signal. (3) A moving dark bar with 5° width moving in 12 possible directions. The sequence of directions or orientations was pseudorandomized.

For the wide-field imaging setup, a large LED screen was placed 15 cm away from the mouse’s eye covering a visual field of 145° x 121°. The center of the monitor was at 25° elevation and either 0° azimuth (covering both eyes), 45° azimuth (mostly right eye) or −45° azimuth (mostly left eye). To avoid excessive fisheye distortion in these wide-field experiments, we applied a spherical correction to the images on the monitor ^35^.

### Viral injection

We injected adeno-associated virus (AAV) expressing non-floxed GCaMP6 (AAV2/1.hSyn1.GCaMP6f.WPRE.SV40) into the SC of wild-type mice or the SC of partial cortex mutant mice. After 2–3 weeks, we implanted a cranial window coupled to a transparent silicone plug that rested on the surface of the SC and exposed its caudomedial portion. This portion of the SC corresponds to the up-temporal part of the visual field. The optics remained clear for several months, which enabled long-term monitoring of the same neurons. Two-photon microscopy was applied to image calcium signals in the SC of head-fixed awake mice 3 weeks to 2 months after viral injection.

### 2-photon calcium imaging and analysis

For imaging experiments, the animal was fitted with a head bar, and head-fixed while resting on a rotating treadmill. The animal was awake and free to move on the treadmill, but not engaged in any conditioned behavior. Two-photon imaging was performed on a custom-built microscope with a 16 ×, 0.8 NA, 3mm WD objective (Nikon) and controlled by custom software written in Labview (National Instruments). A Ti:Sapphire laser with mode-locking technique (Spectra-Physics Mai Tai HP DeepSee) was scanned by galvanometers (Cambridge). GCaMP6f was excited at 920 nm and laser power at the sample plane was typically 20-80 mW. A 600 μm × 600 μm field of view was scanned at 4.8 Hz as a series of 250 pixel × 250 pixel images and the imaging depth was up to 350 μm. Emitted light was collected with a T600/200dcrb dichroic (Chroma), passed through a HQ575/250m-2p bandpass filter (Chroma), and detected by a photomultiplier tube (R3896, Hamamatsu). Artifacts of the strobed stimulus were eliminated by discarding 8 pixels on either end of each line.

Brain motion during imaging was corrected using SIMA^36^ or NoRMCorre^37^. Regions of interest (ROIs) were drawn manually using Cell Magic Wand Tool (ImageJ) and fitted with an ellipse in Matlab. Fluorescence traces of each ROI were extracted after estimating and removing contamination from surrounding neuropil signals as described previously^15, 38^. Slow baseline fluctuations were removed by subtracting the eighth percentile value from a 15-s window centered on each frame^39^.

Two criteria were applied to interpret ROIs as neurons: 1) The size of ROI was limited to 10 – 20 µm to match the size of a neuron; 2) ROIs with a signal-to-noise ratio (SNR) smaller than 0.4 were discarded from further analysis^40^.

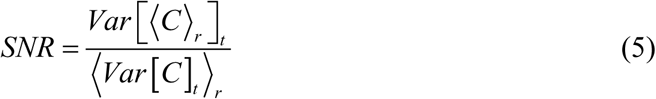

Where *C* is the *T* × *R* response matrix (time samples × stimulus repetitions), 〈 〉 is the mean and *Var* [] is the variance.

For any given stimulus, the response of a neuron was defined by the fluorescence trace in its ROI during the stimulus period

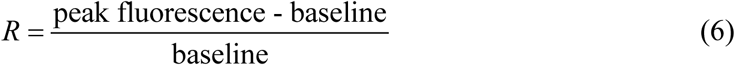

where the baseline is derived from a 0.5-s period prior to stimulation.

To quantify the tuning of a neuron to motion direction, we computed the preferred direction as the argument of the response-weighted vector sum of all directions:

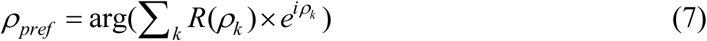

where *R*(*ρ_k_*) is the response amplitude in the *k*th direction *ρ_k_*. The direction selectivity index (DSI) was calculated as the normalized amplitude of the sum:

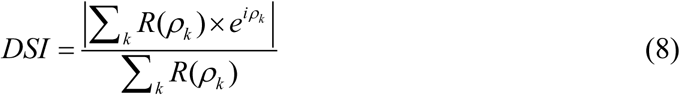

Similarly, for orientation tuning, the preferred orientation was calculated as the argument of the response-weighted vector sum of all orientations:

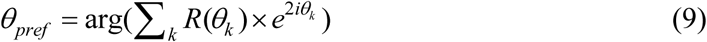

Where *R*(*θ_k_*) is the response amplitude at the *k*th orientation *θ_k_*. The orientation selectivity index (OSI) was calculated as the normalized amplitude of the sum:

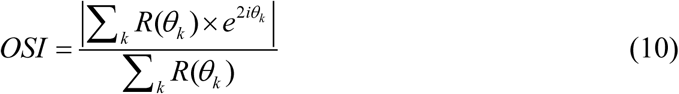

We assessed the statistical significance of DSI and OSI using a permutation test^41^: Surrogate trials were generated by randomly shuffling the trial labels and recomputing the DSI or OSI. This procedure was repeated 1000 times to generate a null distribution. Neurons were classified as direction-selective if *DSI* > 0.1 and *p* < 0.05, and as orientation-selective if *OSI* > 0.1 and *p* < 0.05.

The direction map was generated by linearly interpolating the single-cell-resolution data and then applying a mean filter over a 200-µm square region. To calculate the gradient of the map, the map of the preferred direction was fitted by:

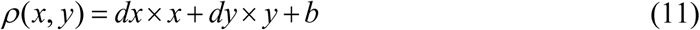

where *x* and *y* are the anatomical coordinates. The gradient was calculated as:

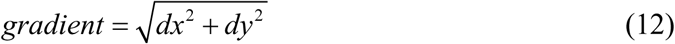

The artificial random arrangement of preferred direction (Fig. 2m) was simulated as 26 × 26 neurons, each of which prefers a random direction, evenly distributed on a 500 µm × 500 µm anatomical space.

To find the optimal number of Gaussians to fit the preferred direction distribution (Fig. 2n), we evaluated the Bayesian information criterion (BIC)^42^:

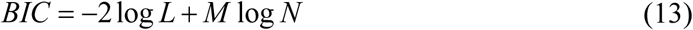

where *L* is the maximized likelihood, *M* is the number of parameters in the model, and *N* is the number of neurons.

The center of the receptive field (RF) of each neuron was calculated based on the moving-bar responses. For each direction of motion, the time of peak response depends on the location of the RF and the neuron’s response latency. By evaluating responses to all directions, one can extract the response latency and determine the best-fit location of the RF.

### Wide-field imaging and analysis

The wide-field microscope was built with a 50-mm / 105-mm tandem lens^43^. Illumination was from a blue LED with an excitation filter (469 nm ± 35 nm) and the camera used an emission filter (525 nm ± 39 nm). Images were acquired using a CMOS camera (Basler ace acA2000-165µm NIR USB 3.0 Camera) at an acquisition rate of 10 Hz. The camera covers an area of 5.36 mm × 2.85 mm at a spatial resolution of 2.63 µm. Each acquired image was convolved with a normalized box filter and down-sampled to 1/10 of the original size. An ROI was drawn manually over the SC to exclude signals from the surrounding areas. For each pixel in the image, the response was defined as described above for ROIs in 2-photon experiments. The RF center of each pixel was calculated from moving bar responses as described above.

To measure the size of direction-tuned patches in a direction map, we calculated the normalized 2-D autocorrelation of the map. Similarly to evaluate the correspondence between maps on the left and right SC or across different mice, we calculated the normalized 2-D cross-correlation of two maps. Suppose a direction map is given by 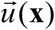, where at each location **x** in the visual field 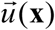 is a vector that points in the preferred direction *ρ* with length proportional to the DSI. For two such direction maps 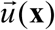 and 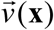 we define the cosine correlation as

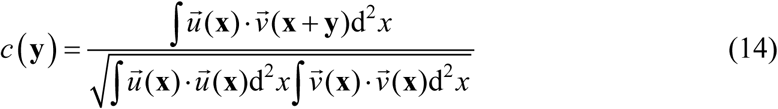

For every possible shift vector **y**, this computes the normalized dot product of one map with the shifted version of the other map. We averaged this correlation map over all shift vectors of the same distance |*y*| = *r* to obtain a radial profile *C*(*r*). The full width at half maximum (FWHM) of the correlation map (Fig. 7b) was defined as twice the radius where this profile dropped to half of its maximum.

### Statistical methods

No statistical method was used to predetermine the sample size. A permutation test was used to assess the statistical significance of DSI and OSI as described above. The Shapiro-Wilk test was applied to assess whether data were normally distributed. In the case of a normal distribution, a *t*-test was applied; otherwise, a nonparametric test (Mann–Whitney U-test or One-sample Wilcoxon signed-rank test) was applied.

## Acknowledgements

M.M was supported by a grant from NIH (R01 NS111477). Y.L.was supported by NEI K99EY028640 and a Helen Hay Whitney Postdoctoral Fellowship.

## Author Contributions

Y.L. designed the study, performed all experiments, interpreted results, and wrote the manuscript. Z.T. provided partial cortex mutant mice and validated their brain anatomy. M.M. helped design the study, interpret results, and write the manuscript.

## Data and Code availability statement

Data and code used in generating figures will be available on Github following acceptance of the manuscript.

## Figure Legends

**Supplementary Figure 1.**
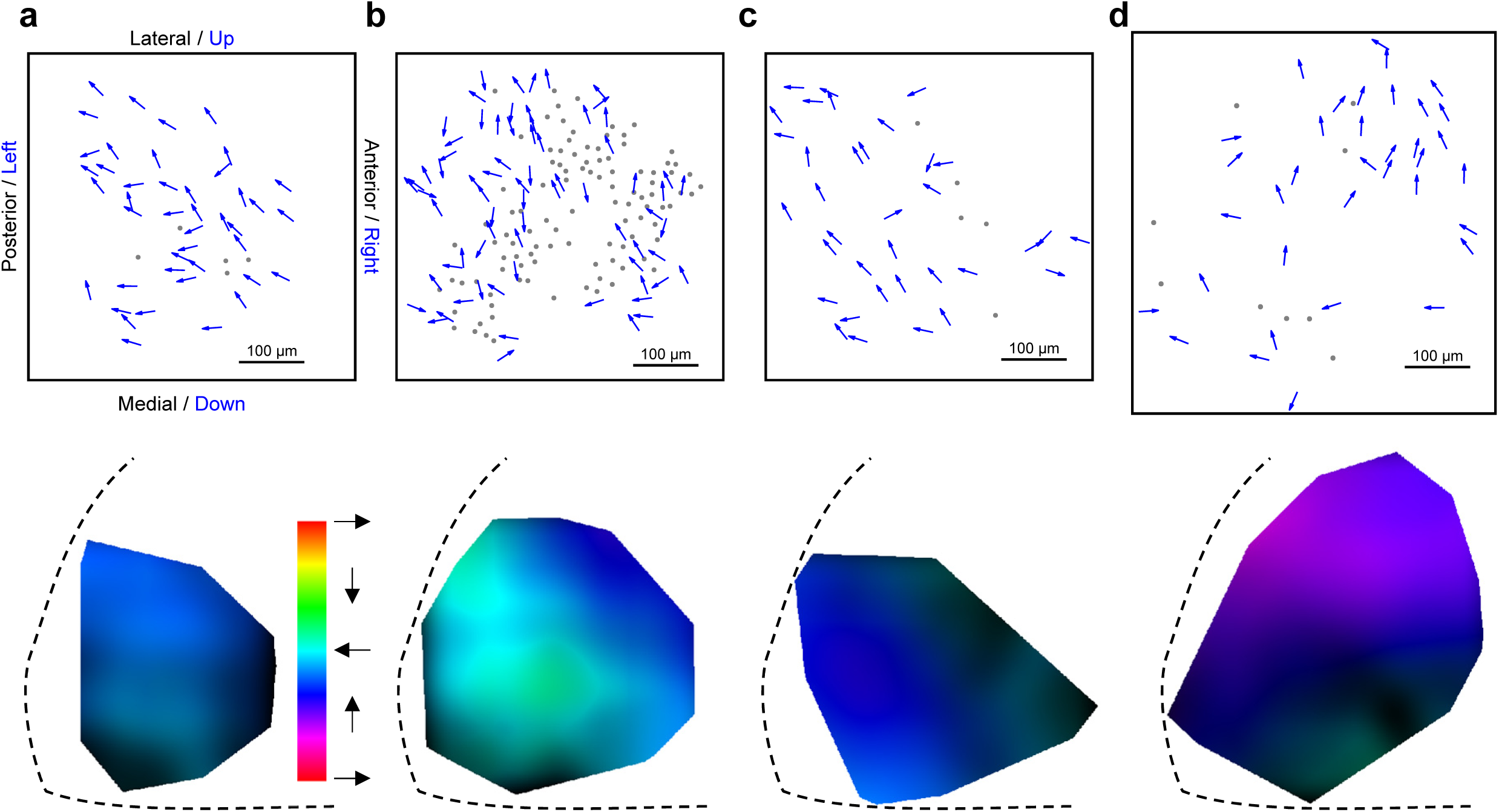
Additional examples of arrow plots and smooth maps of preferred directions. Data from 4 animals stimulated with moving dots, presented as in Figs 2a and 2c.

**Supplementary Figure 2.**
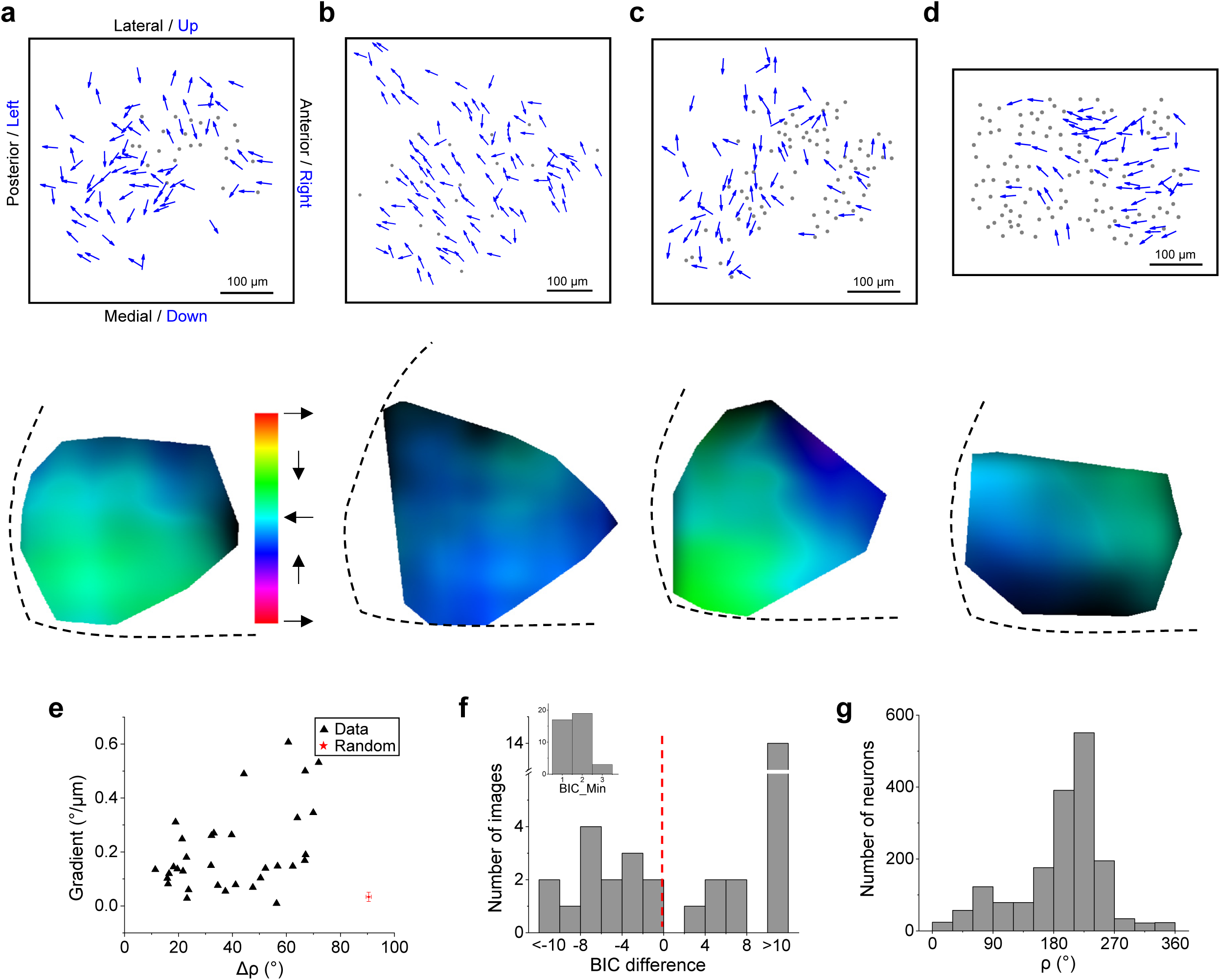
Organization of motion direction measured by moving bars. (**a**, **b**, **c**, **d**) Data from 4 animals stimulated with moving bars, presented as in Figs 2a and 2c. **e**, Gradients of the map plotted against the absolute difference in preferred directions for pairs within 50 µm (33 images), displayed as in Fig. 2m. **f**, Analysis of bimodality in the distributions of preferred direction (33 images), displayed as in Fig. 2n. **g**, Histograms of preferred direction for all direction-selective neurons (1754 cells).

**Supplementary Figure 3.**
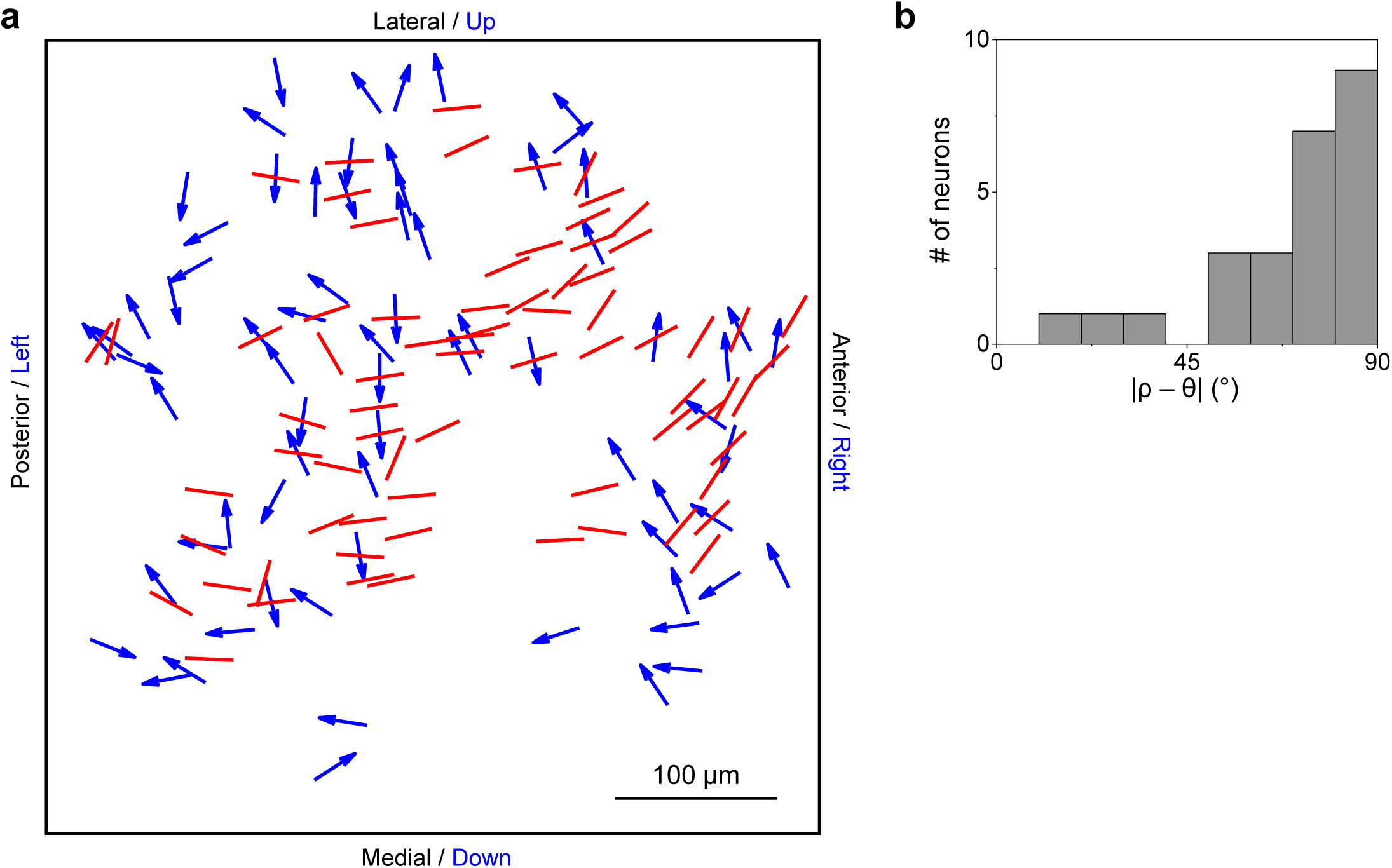
Another sample showing the orthogonal relationship between preferred direction and orientation. **a.** Overlay of arrow plots for direction-selective neurons and line plots for orientation-selective neurons. **b**, Histogram of the absolute difference between preferred direction and orientation (25 cells).

**Supplementary Figure 4.**
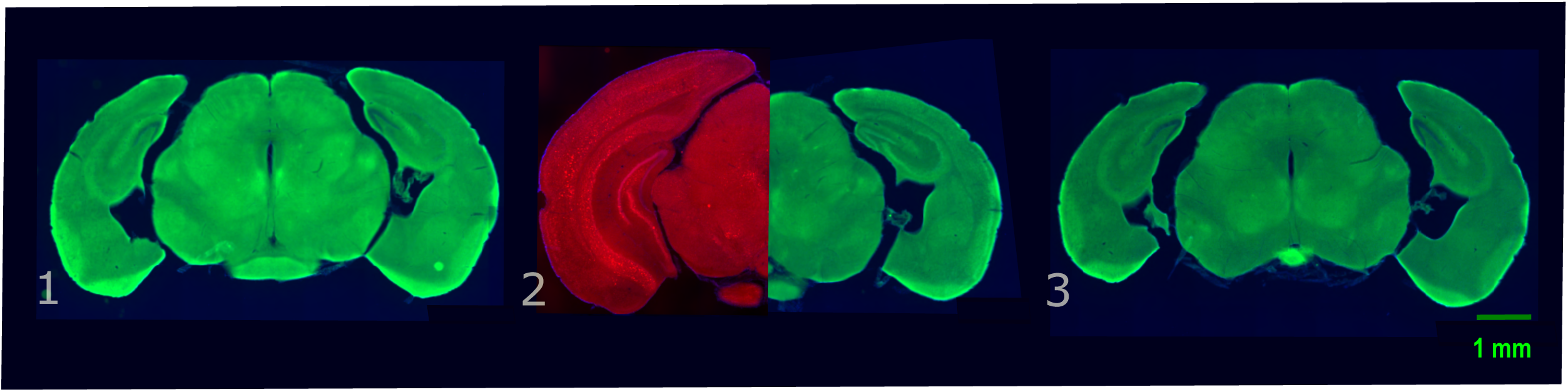
Brain anatomy of a mutant mouse (Emx1-Cre:Pals1^flox/wt^) with partially developed neocortex. Series of coronal sections (100 µm thick), labeled from anterior (1) to posterior (3). The left half of panel 2 shows the corresponding section from a wild-type mouse brain. Autofluorescence (green, red).

**Supplementary Figure 5.**
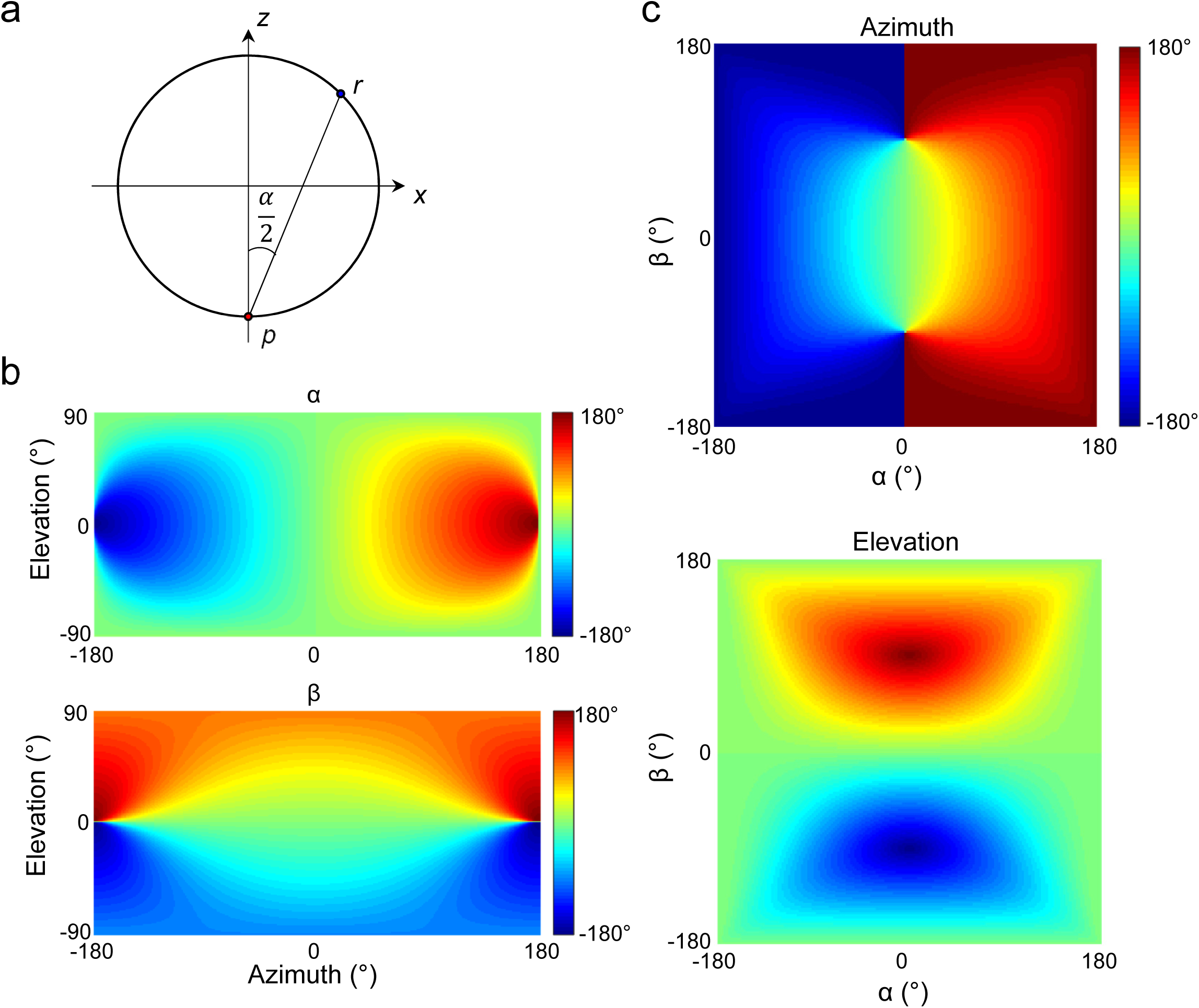
Introduction of unipolar coordinates. **a,** Schematic of the definition of unipolar coordinates (see **Methods**). **b and c**, Relationship between spherical coordinates and unipolar coordinates.

**Supplementary Figure 6.**
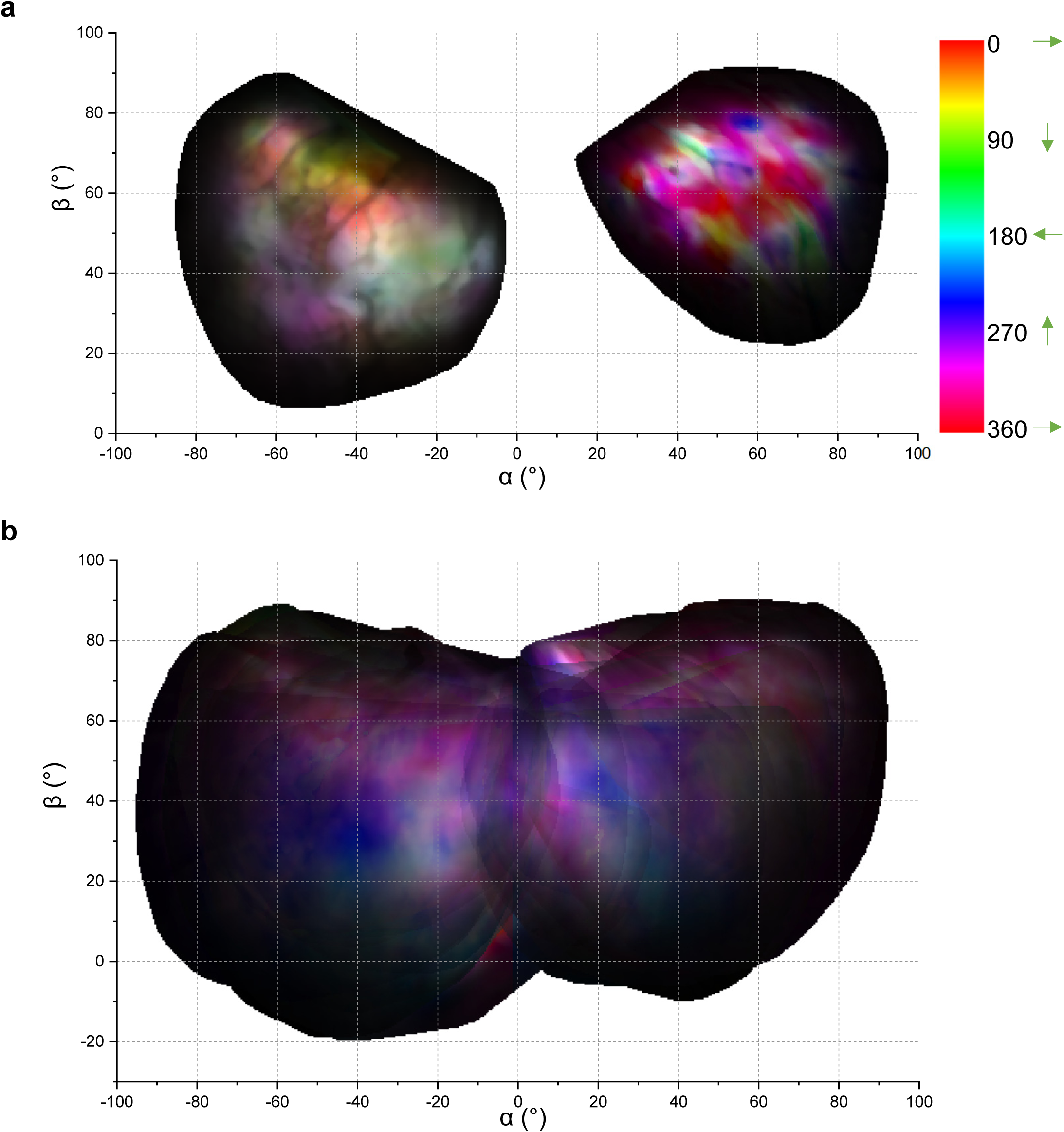
Direction maps including both sides of the SC. **a,** A sample direction map plotted in visual space with unipolar coordinates. Hue codes the preferred direction. Value codes the amplitude of responses. Saturation codes the DSI. The horizontal component of the direction in the left field has been flipped to better reveal left-right symmetry. **b**, Average direction map combined over 9 animals.

**Supplementary Figure 7.**
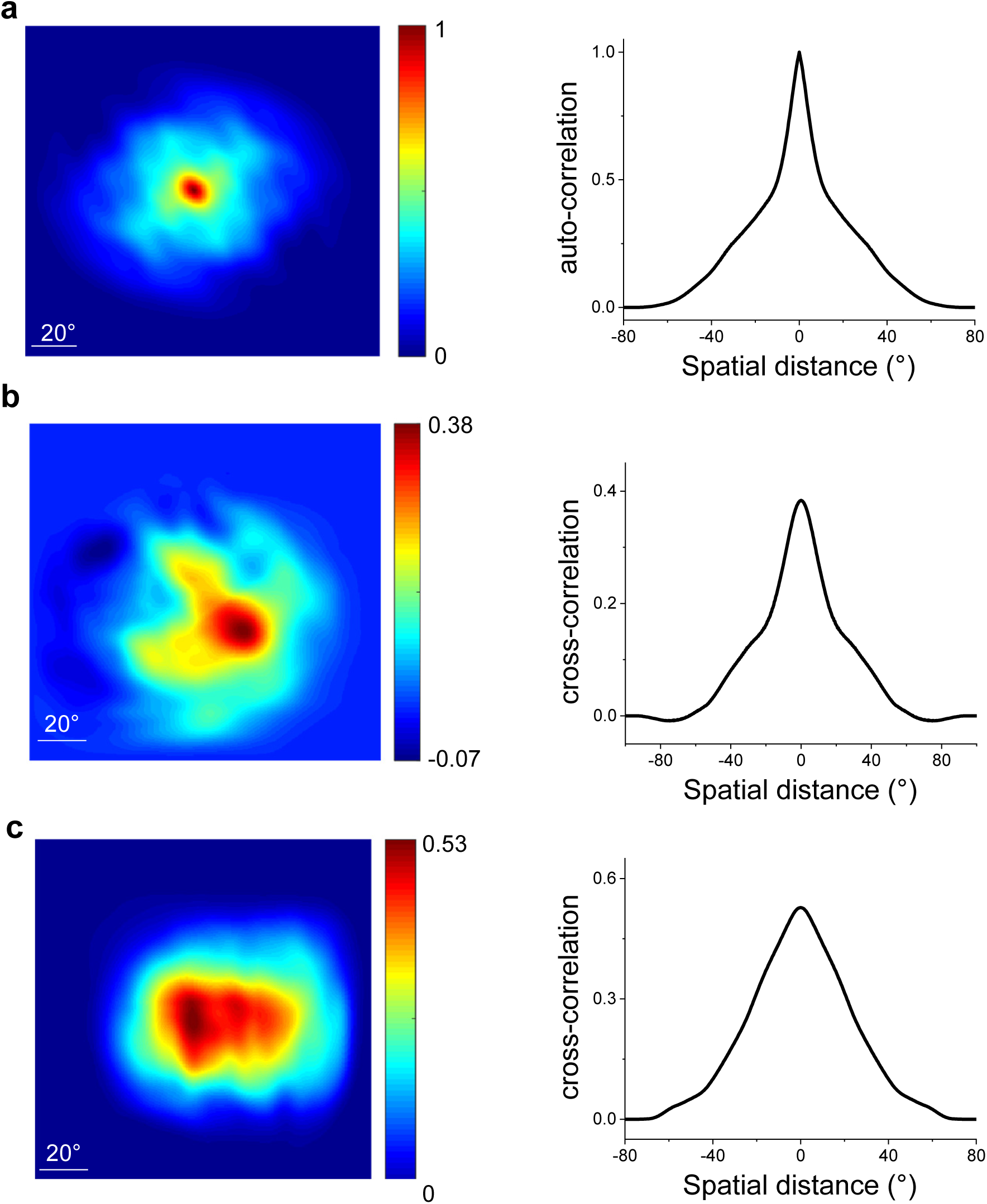
Autocorrelation and cross-correlation of global direction maps. **a,** Left, the autocorrelation of a sample direction map. Right, the corresponding radial profile. **b,** Left, the cross-correlation between direction maps of the left and right SCs of an example animal. Right, the corresponding radial profile, centered on the peak of the correlation map. **c,** Left, the cross-correlation between direction maps of the same-side SCs of two different animals. Right, the corresponding radial profile.

**Supplementary Figure 8.**
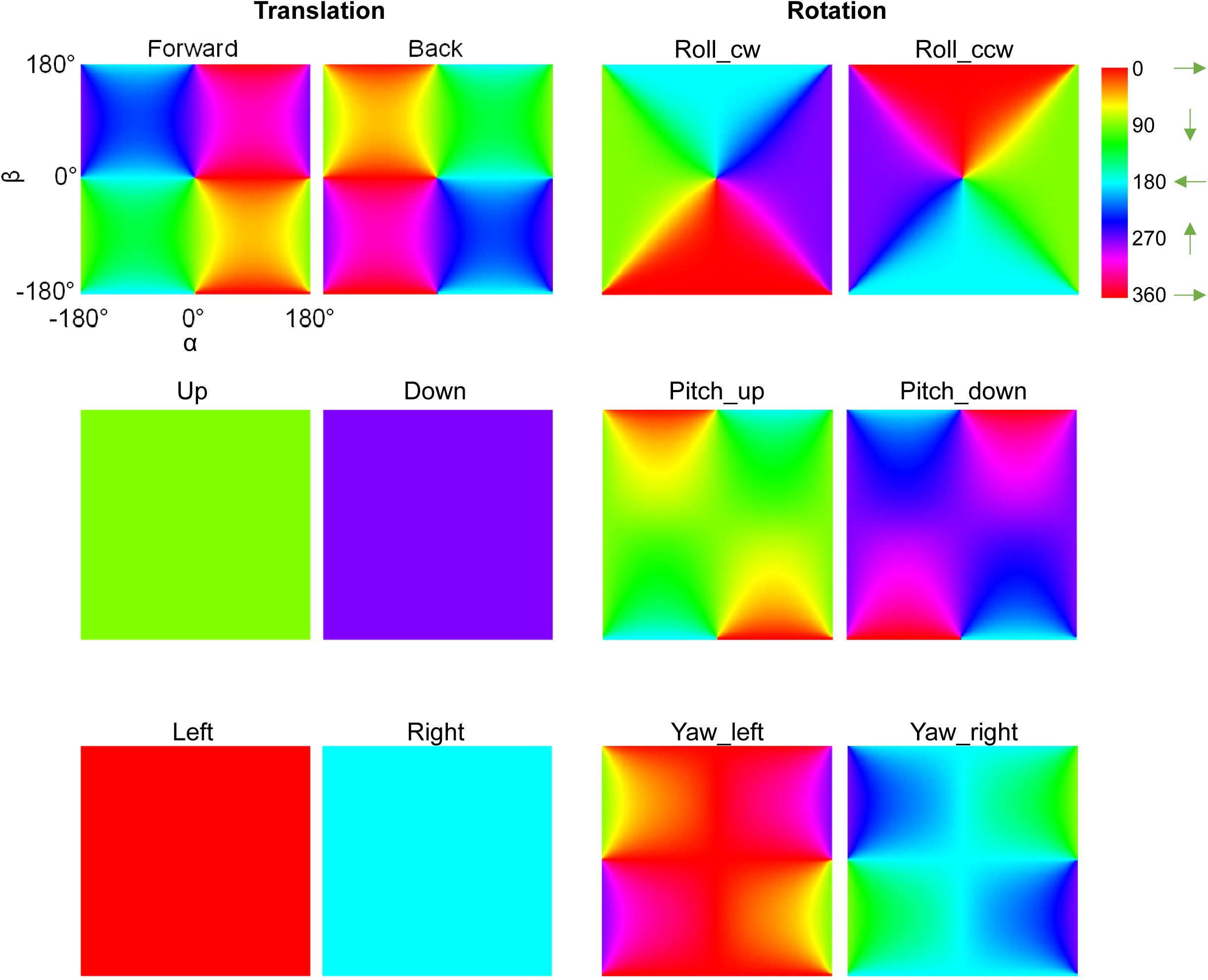
Motion maps corresponding to optic flows from self-motion. For each of the 6 basic modes of translation and rotation, this plots the direction of the optic flow vectors in the visual field using unipolar coordinates. Note these graphs assume that the axis of forward translation and roll rotation equals the lambda-bregma axis. In actuality, a freely moving mouse may carry the head inclined downward during ambulation^44^ or upward during rearing.

**Supplementary Figure 9.**
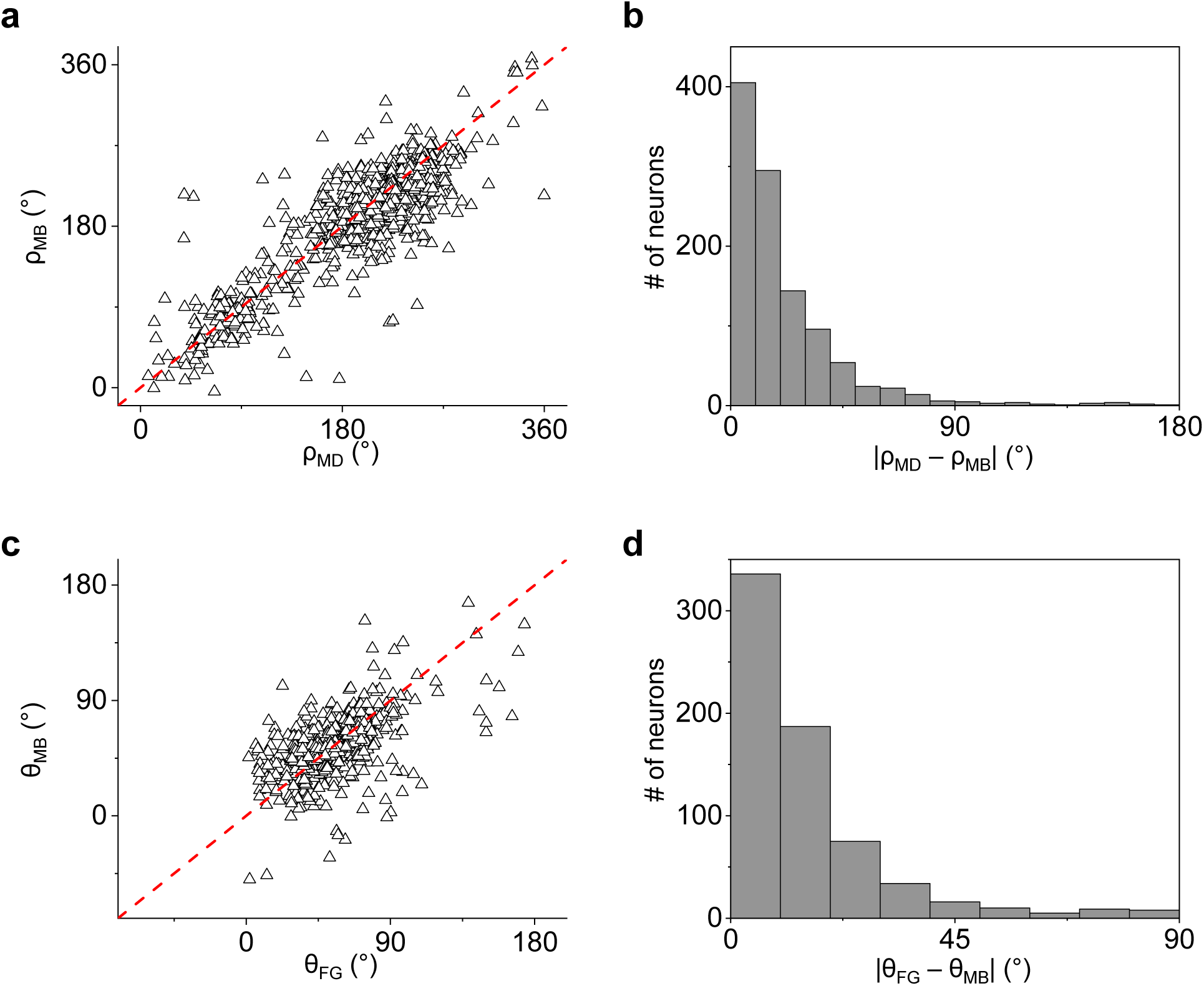
Comparison of preferred direction or orientation measured by different stimuli. **a,** Preferred direction measured by moving bars plotted against that by moving dots (1085 neurons from 14 animals). **b,** Histogram of difference between preferred directions measured by these two stimuli. **c,** Preferred orientation measured by moving bars plotted against that by flashed gratings (680 neurons from 14 animals). **d,** Histogram of difference between preferred orientations measured by these two stimuli.

